# Machine learning predicts immunoglobulin light chain toxicity through somatic mutations

**DOI:** 10.1101/849901

**Authors:** Maura Garofalo, Luca Piccoli, Margherita Romeo, Maria Monica Barzago, Sara Ravasio, Mathilde Foglierini, Milos Matkovic, Jacopo Sgrignani, Raoul De Gasparo, Marco Prunotto, Luca Varani, Luisa Diomede, Olivier Michielin, Antonio Lanzavecchia, Andrea Cavalli

## Abstract

In systemic light chain amyloidosis (AL), pathogenic monoclonal immunoglobulin light chains (LCs) form toxic aggregates and amyloid fibrils in target organs. Prompt diagnosis is crucial to avoid permanent organ damage. However, delays in diagnosis are common, with a consequent poor patient’s prognosis, as symptoms usually appear only after strong organ involvement. Here, we present LICTOR, a machine learning approach predicting LC toxicity in AL, based on the distribution of somatic mutations acquired during clonal selection. LICTOR achieved a specificity and a sensitivity of 0.82 and 0.76, respectively, with an area under the receiver operating characteristic curve (AUC) of 0.87. Tested on an independent set of 12 LCs sequences with known clinical phenotypes, LICTOR achieved a prediction accuracy of 83%. Furthermore, we were able to abolish the toxic phenotype of an LC by *in silico* reverting two germline-specific somatic mutations identified by LICTOR and by experimentally assessing the loss of *in vivo* toxicity in a *Caenorhabditis elegans* model. Therefore, LICTOR represents a promising strategy for AL diagnosis and reducing high mortality rates in AL.

## Introduction

Systemic light chain amyloidosis (AL) is a monoclonal gammopathy characterized by the abnormal proliferation of a plasma cell clone producing large amounts of pathogenic immunoglobulin free light chains (LCs)^1^. LCs, mainly secreted as homodimers^2^, misfold forming toxic species and amyloid fibrils which accumulate in target organs and lead to fatal organ dysfunction and death^1^. Although LCs deposition can occur in any organ except the brain, the kidney and heart are the most affected sites, with the latter bearing the worst prognosis. Symptoms of AL are non-specific and usually reflect advanced organ involvement. Therefore, an early diagnosis is essential to avoid irreversible organ damage. However, the complexity of the disease and its vague symptoms make a timely diagnosis of AL extremely challenging^3,4^.

Pre-existing monoclonal gammopathy of undetermined significance (MGUS) is a known risk factor for developing AL, with 9% of MGUS patients progressing to AL^5–7^. However, early diagnosis is still difficult since reliable diagnostic tests predicting whether MGUS patients are likely to develop AL are currently lacking^7,8^. Predicting the onset of AL is highly challenging, as each patient carries a different pathogenic LC sequence resulting from a unique rearrangement of variable (V) and joining (J) immunoglobulin genes and a unique set of somatic mutations (SMs) acquired during B cell affinity maturation^9^ (Fig. 1a). Therefore, the development of a specific prediction tool represents a crucial step to anticipate AL diagnosis and improve patients’ prognosis.

**Fig. 1.**
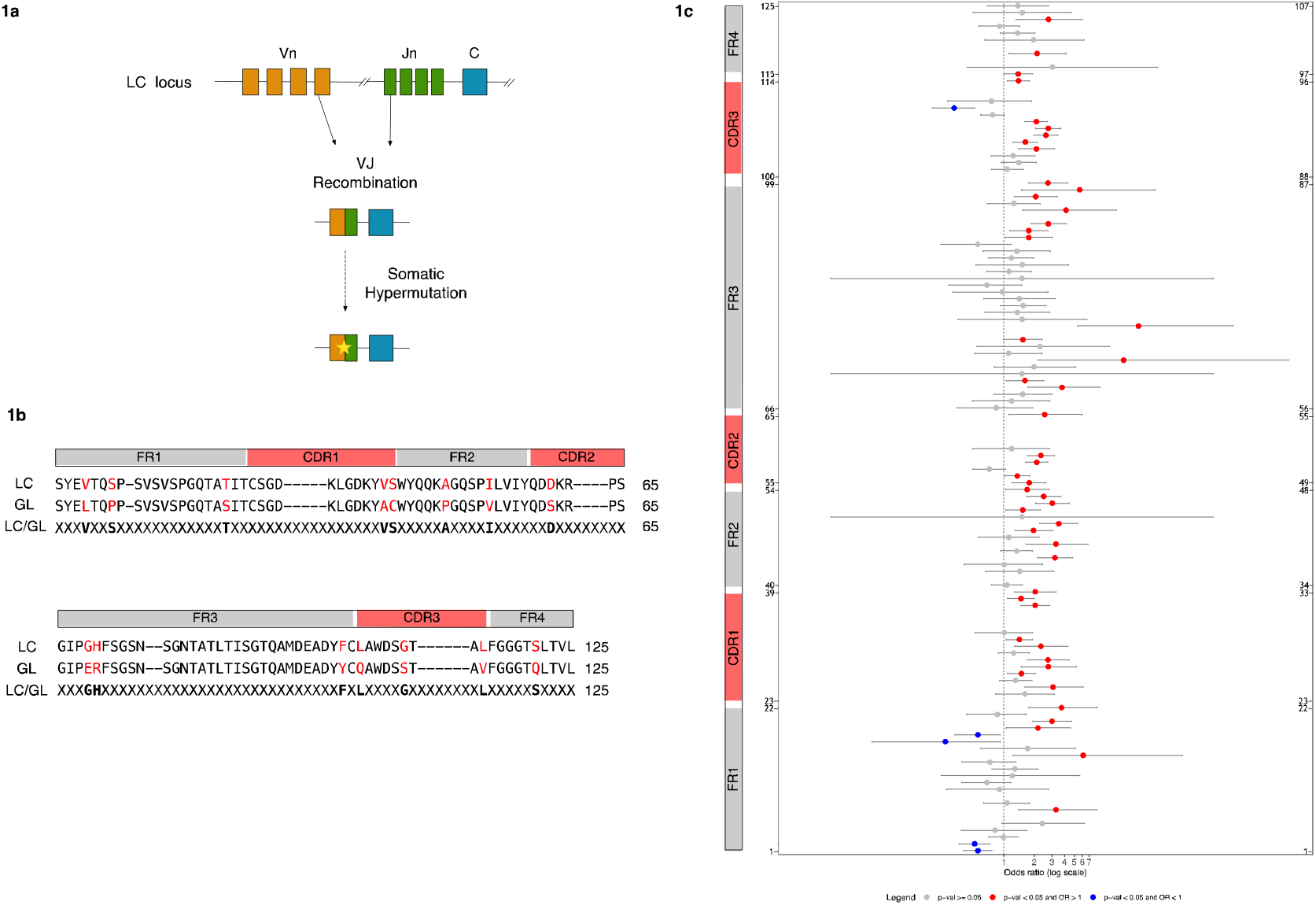
The presence of somatic mutations differentiates toxic and non-toxic LC sequences. **a**, Schematic representation of the generation of LC diversity through the processes of VJ recombination and somatic hypermutation. **b**, Alignment of an LC sequence with the corresponding germline sequence (GL) according to Kabat-Chothia scheme using a progressive enumeration for a total of 125 positions (Methods). Structural elements of immunoglobulin light chains are depicted on top of the sequences (FR1= framework 1, CDR1= complementary determining region 1, FR2= framework 2, CDR2= complementary determining region 2, FR3= framework 3, CDR3= complementary determining region 3, FR4= framework 4). Residues in red depict somatic mutations. The third line shows the encoding scheme used by the classifier with somatic mutations (displayed in bold) and unmutated positions represented by an ‘X’. **c**, Odds Ratio (OR) for all 125 positions of the LC sequences according to our sequential numbering scheme (y-axis). The corresponding Kabat-Chothia enumeration is reported on the right. Structural elements of immunoglobulin light chains are shown on the left. ORs for positions with no statistically significant difference between *tox* and *nox* sequences (p ≥ 0.05) are represented as grey dots. Positions with statistically significant differences (p < 0.05) are depicted as either red (OR>1) or blue (OR<1) dots. Grey horizontal error bars show the OR 0.95 confidence interval.

Machine learning techniques are becoming very prominent in different areas of science and are also gaining acceptance in medicine. Indeed, machine learning has been used in different areas of medicine, such as diagnosis^10–12^, prognosis^13,14^, drug discovery^15,16^ and drug sensitivity prediction^17–19^. In these approaches, machines learn information from data without being explicitly programmed and simulate human intelligence to make predictions^20^. The high diversity of LC sequences accountable for AL development and the possibility of accessing databases of pathogenic and non-pathogenic LC sequences prompted us to use a machine learning-based strategy to devise a predictor of LC toxicity in AL named LICTOR (λ-**LI**ght-**C**hain **TO**xicity predicto**R**).

LICTOR uses SMs as predictor variables based on the hypothesis that SMs are the main LC toxicity-discriminating factors, achieving a specificity and a sensitivity of 0.82 and 0.76, respectively, with an AUC of 0.87. LICTOR performance was assessed with an independent set of LCs with a known clinical phenotype but not used in the training. Furthermore, to experimentally validate LICTOR, we used our predictor to abolish the pathological phenotype of a cardiotoxic LC and verified the outcome with a *Caenorhabditis elegans*-based assay^21,22^. This assay takes advantage of the evolutionary relationship between the nematode pharynx and the vertebrate heart, evaluating the reduction of the *C. elegans* pharyngeal pumping rate after administration of cardiotoxic LCs as a measure of proteotoxicity^21,22^. Both experiments recapitulated LICTOR’s predictions.

Taken together, these results confirm that LICTOR provides insights into specific features differentiating toxic and non-toxic LCs. Therefore, it may represent a powerful tool to improve AL diagnosis and unveil a novel strategy for patient treatment through personalized medicine.

## Results

### Somatic mutations are key LC toxicity-discriminating factors

To investigate the role of SMs in the generation of toxic LCs and validate their use as predictor variables in a LC toxicity predictor, we collected a database of 1,075 λ LC sequences. The database included 428 “toxic” sequences (i.e., LCs responsible for the formation of toxic aggregates an AL development) extracted from AL patients (hereafter referred to as *tox*) and 647 “non-toxic” LCs (*nox*) comprising sequences from healthy donor repertoires, other autoimmune diseases or cancer, obtained from ALBase^23^ (428 *tox*, 590 *nox*) and an in-house LCs database not related to AL (57 *nox)*. We restricted our analysis to λ LCs since this isotype is more prevalent than the kappa (κ) isotype in AL patients (λ/κ=3:1 compared to that of healthy individuals, λ/κ=1:2)^24^. To identify SMs, all LCs were aligned to the corresponding germline (GL) sequence obtained using the IMGT database^25^. LCs were then numbered according to the Kabat-Chothia scheme (using a progressive enumeration from 1 to 125), allowing the structural comparison of LCs with different sequence lengths (Methods and Fig. 1b). Next, we counted the number of mutated (M) and non-mutated (NM) residues at each position *i* in *tox* and *nox* sequences (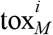 and 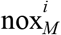, 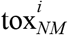 and 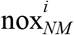, respectively) and used Fisher’s exact test^26^ to assess whether the frequencies of mutations 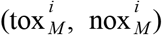 and non-mutations 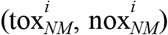 were significantly (p < 0.05) different. Finally, the odds ratio (OR)^26^ was used to assess the association strength between mutations and toxicity at each position *i* in *tox* and *nox* sequences (Methods and Fig. 1c). Interestingly, 48 of 53 positions with a statistically significant difference (p < 0.05) between the two groups (Fig. 1c) showed a higher rate of mutation in the *tox* group (OR >1), while only 5 positions reported a higher mutation rate in the *nox* group (OR <1). To exclude possible bias induced by the use of a group of *nox* sequences having an artificially low level of SMs, we randomly selected 1,000 LC sequences from a healthy donor repertoire (*hdnox*)^27^ and compared the probability distributions of the number of SMs (PDSM) between the three groups. We observed similar PDSM between the *nox* and *hdnox* groups, while the PDSM of *tox* and *hdnox*, as well as *tox* and *nox*, were significantly different. This result supports *nox* sequences as a bona fide group of LCs (Supplementary Figure 1). Overall, these findings suggest that SMs are key determinants of the toxicity of LCs and, thus, can be used as predictor variables to develop LC toxicity prediction tools.

### Prediction of LC toxicity using machine learning

The previous findings prompted us to use SMs as features to develop a machine learning approach automatically classifying LCs as either toxic or non-toxic in AL. To this end, we combined the information from SMs with knowledge of the 3D structure of LCs homodimers^28,29^ to create three families of predictor variables used in the training of machine learning algorithms. The first family, termed AMP (Amino acid at each Mutated Position), highlights sequence features, identifying the presence or absence of an SM at each position of the LCs sequences. The second family, termed MAP (Monomeric Amino acid Pairs), identifies the presence or absence of mutations in residues in close contact in the LC monomeric 3D structure (distance <7.5 Å). Finally, the third family, named DAP (Dimeric Amino acid Pairs), identifies the presence or absence of mutations at positions in close contact but belonging to different chains. Next, four machine learning algorithms (Bayesian network, logistic regression, J48 and random forest)^30^ were evaluated for their ability to solve the classification problem, using our database as input. To assess the importance of the different classes of predictor variables, we performed 28 prediction experiments including all possible combinations of AMP, MAP and DAP families. In addition, to avoid unbalanced class problems, i.e., the tendency of machine learning algorithms to assign sequences to the largest class in the dataset, *nox* in our case, each of the 28 experiments was performed with and without balancing the training set using a SMOTE (Synthetic Minority Over-sampling TEchnique) filter^31^. The assessment of algorithms and predictor variable combinations was performed using 10-fold cross-validation to avoid overfitting. We found that for all tested machine learning algorithms, the best combination of predictor variable families provided an area under the receiver operating characteristic curve (AUC) that substantially differed from that of a random classifier (0.50), with random forest being the best classifier (0.87) and J48 being the worst (0.75) (Fig. 2a and Supplementary Table 1). Furthermore, all four classifiers relied on the AMP family to predict LC toxicity, while only random forest used all three families of predictor variables (AMP+MAP+DAP) in its best configuration. Overall, these findings highlight the importance of the structural context of somatic mutations in defining the toxicity of an LC and identify random forest using AMP, MAP, and DAP as the best approach in our case. For this reason, we used random forest in our implementation of LICTOR.

**Fig. 2.**
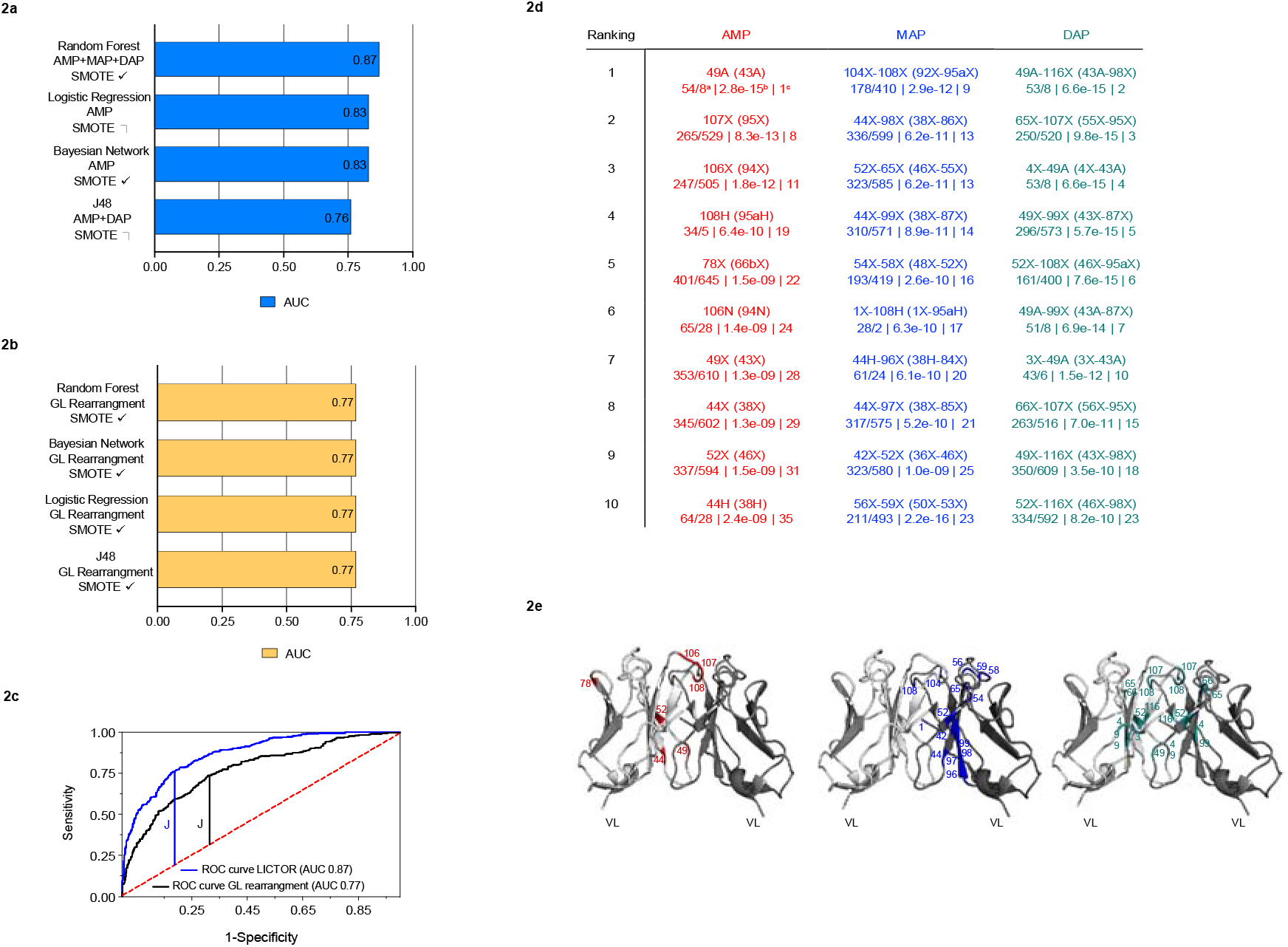
Machine learning predicts toxic and non-toxic sequences and identifies key features of toxicity. **a**, AUC of the best configuration for each of the considered machine learners (blue bars). Different combinations of three families of predictor variables were tested, with (@) or without (@) the SMOTE balancing technique. **b**, The yellow bars show the best AUC value obtained by each machine learner using only the LC germline VJ rearrangements as predictor variables. **c**, ROC curve for LICTOR (i.e., random forest using AMP + MAP + DAP) compared with a predictor (random forest) using only the LC germline VJ rearrangements as predictor variables. **d**, Top 10 features of each family ranked by information gain. Each feature is enumerated according to our sequential numbering scheme, while the corresponding Kabat-Chothia enumeration for each feature is reported in parenthesis. Kabat-Chothia insertions are reported with lowercase letters. Below each predictor variable are shown the occurrence in tox/nox sequences (a), the p-value (b) and the feature selection general ranking (c) (red= AMP features, blue= MAP features, and green=DAP features). **e**, Mapping of the top 10 features of each family on the variable domains of an LC homodimeric structure (PDB ID: 2OLD, represented in white and grey in cartoon). AMP features are shown in red in the left image, MAP features in blue in the middle image, and DAP features in green in the right image. The colour code used in the table to represent the three feature families is maintained in their structural representation in **d**.

### Validation of LICTOR

Next, we sought to validate the prediction accuracy of LICTOR with a set of sequences with known clinical phenotypes but not present in the training set (*valset*).^29^ The *valset* (Supplementary Table 2) comprised a total of 12 LCs, including 7 sequences associated with AL with cardiac involvement (H3, H6, H7, H9, H15, H16 and H18) and 5 from multiple myeloma patients (M2, M7, M8, M9 and M10). LICTOR was able to correctly classify 10 (6 *tox* and 4 *nox*) of 12 LCs as either toxic or non-toxic (Supplementary Table 2 and Fig. 3a). The probability of achieving a similar accuracy with a random classifier was 0.016, strengthening the argument that LICTOR is a robust and accurate tool to predict the clinical toxicity of previously unseen LCs.

**Fig. 3.**
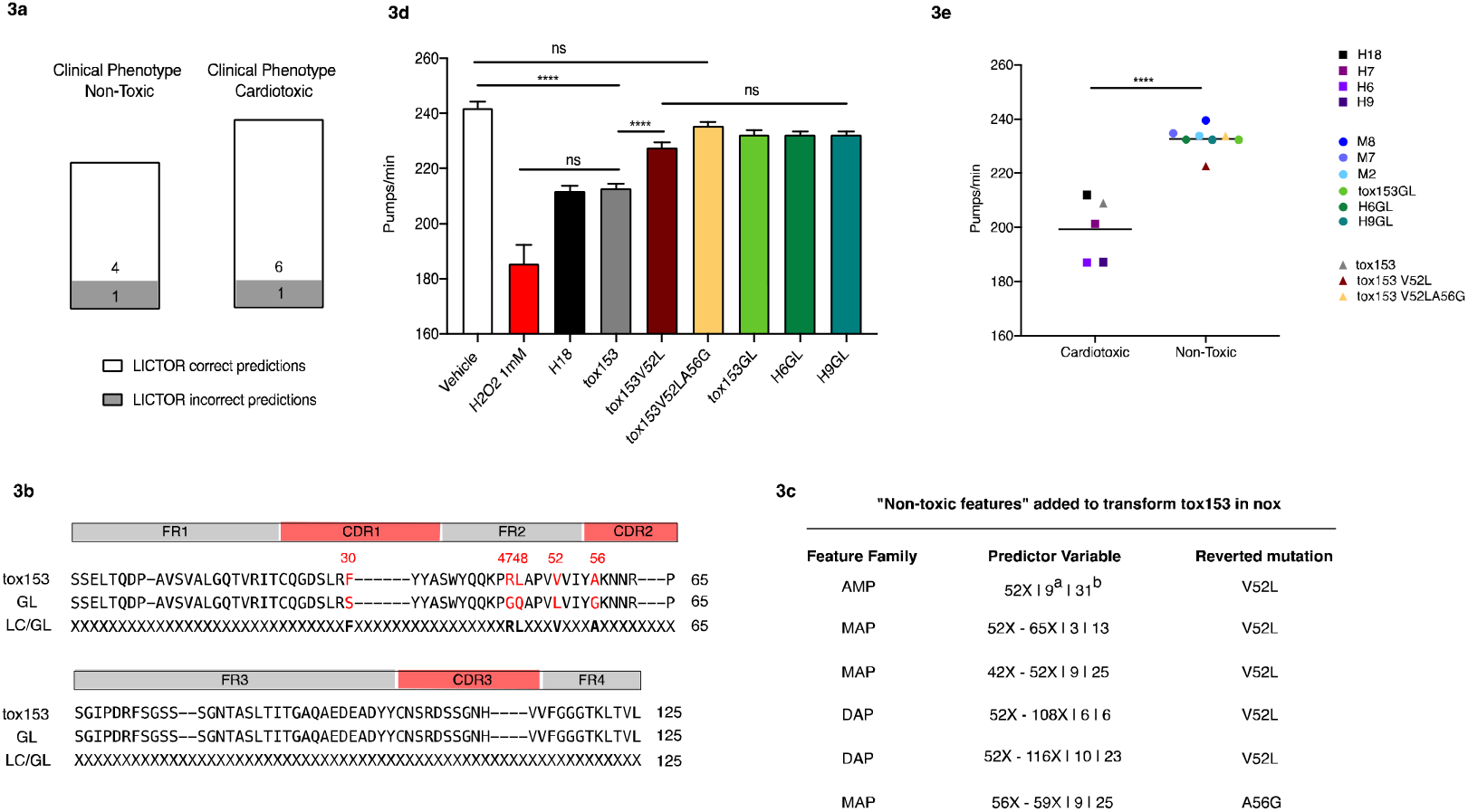
LICTOR accurately predicts the LC toxicity of sequences absent from the training set and is able to revert the pathological phenotype of a cardiotoxic LC. **a**, LICTOR predictions based on an independent set of LCs, i.e., not present in the training set. Toxic LCs are from patients affected by AL with cardiac involvement, while non-toxic LCs are from patients with multiple myeloma (see also Supplementary Table 2). Predictions are divided according to the clinical phenotype. White bars represent correct LICTOR predictions, while grey bars represent incorrect predictions. **b**, Sequence of a cardiotoxic LC (tox153) used to neutralize the toxic phenotype using LICTOR and the non-toxic features unveiled by feature selection. Tox153 is aligned with the corresponding germline (GL) sequence, and the third line shows the difference in somatic mutations between the LC and the GL sequence (LC/GL) with somatic mutations (displayed in bold) and unmutated positions represented by an ‘X’. **c**, The table represents the non-toxic features according to information gain used to revert the toxic phenotype of tox153. For each predictor variable, we also report the ranking in the specific feature family (^a^) and the feature selection general ranking (^b^) **d**, Proteotoxic effect of tox153 protein, of the two mutants *in silico* designed by adding non-toxic features (tox153V52L and tox153V52LA56G) and of the tox153 GL protein. The proteotoxic effect of H18 cardiotoxic LC and of the germline proteins H6GL and H9GL are tested as well. Proteins in 10 mM PBS (100 μg/ml) were administered to *C. elegans* (100 μl/100 worms). Vehicle (10 mM PBS) and 1 mM H_2_O_2_ were administered as negative and positive controls, respectively. Pharyngeal activity was determined 24 h after treatment by determining the number of pharyngeal bulb contractions (pumps/min). Data are the mean pumps/min ± SE (n= 30 worms/assay, two assays). **** p < 0.0001 one-way ANOVA, Dunn’s *post hoc* test. **e**, Values of pharyngeal bulb contraction (pumps/min) of some LCs (H6, H7, H9 and M2, M7, M8) listed in Supplementary Table 2 are from AL patients with cardiac involvement (Cardiotoxic) and patients with multiple myeloma (Non-Toxic). These values were previously obtained under the same experimental conditions employed in this study^21,22^. Additionally, values of pharyngeal bulb contraction (pumps/min) of H18, H6GL and H9GL are reported as well. Each square is the mean value for H6, H7, H9 and H18 while dots represent the mean of M2, M7, M8, H6GL and H9GL. Horizontal lines represent the mean of cardiotoxic and non-toxic LCs (**** p < 0.0001, unpaired t-test). The values of pumps/min obtained after the administration of tox153, tox153V52L, and tox153V52LA56G are also plotted (triangles).

To further assess the robustness of LICTOR, we performed two additional tests. In the first test, we predicted the toxicity of 100 randomly selected LCs from the healthy donor repertoire *hdnox* (all absent from the training set). In this case, LICTOR correctly classified 80% of the sequences as non-toxic (Supplementary Table 3), thus confirming a similar level of accuracy as for the training set. In the second test, we assessed LICTOR by further verifying the absence of overfitting. To achieve this, we took all *tox* sequences and randomly labelled half of them as *nox*. Then, we trained a classifier using the 10-fold cross-validation on such a dataset obtaining an AUC of 0.5 (Supplementary Table 4), equivalent to that of a random classifier. The same procedure has been used for the *nox* group (by randomly assigned half of them as *tox* and training a classifier on such a dataset), obtaining also in this case an AUC of 0.5 (Supplementary Table 5). These results further underline that *tox* and *nox* sequences have distinctive features allowing their discrimination.

### LC germline VJ rearrangement information does not improve prediction performance

To further underscore the role of SMs as main discriminants between toxic and non-toxic LCs, we trained the same machine learners employed before, using the LC germline VJ rearrangements as a unique predictor variable, given the well-documented overrepresentation of certain VL germline genes in AL^32–34^. All the resulting *germline-based* classifiers achieved an AUC of 0.77 in their best configuration (Fig. 2b and Supplementary Table 6), a value substantially better than that of a random classifier, although much lower than LICTOR’s score (0.87). Interestingly, adding LC germline VJ rearrangements to LICTOR did not improve its prediction performance (Supplementary Table 7).

Next, we computed the specificity and sensitivity of the two random forest predictors, LICTOR and the *germline-based* predictor, maximizing the Youden index (J)^35^ as a function of the *confidence level* of the random forest predictions, i.e., the probability that a sequence belongs to the predicted phenotype (Fig. 2c). LICTOR achieved a specificity of 0.82 and a sensitivity of 0.76 (J=0.58, threshold=0.46 in identifying *tox*), while the *germline-based* classifier showed a 0.69 specificity and a 0.73 sensitivity (J=0.43, threshold=0.48 in identifying *tox*). Overall, these data suggested that SMs harbour key information that can be used to discriminate between *tox* and *nox*, while LC germline VJ rearrangements do not seem to carry additional information that can improve the prediction performance of LICTOR.

### LICTOR unveils specific features of LC toxicity

To identify the key features leading to LC toxicity in AL, we ranked the predictor variables of LICTOR according to their “information gain”, a value representing the importance of the information carried by each predictor variable for the classification^36^. We found that among the top 10 most important features of the three families of predictor variables, feature 49-A, which denotes an SM to alanine at position 49, obtained the highest score in the AMP family ranking, as well as in the general ranking (Fig. 2d and Supplementary Table 8). Indeed, feature 49-A was present in 54 *tox* sequences but only 8 *nox* sequences. Furthermore, the 49-A mutation, which is located at the dimeric interface of LCs (Fig. 2e), was also ranked among the top 10 features in the DAP family in combination with no substitutions at other residue positions (Fig. 2d). Moreover, among the best-ranked predictor variables of the three families, those describing mutated positions were more frequent in *tox* sequences than in *nox* sequences (Fig. 2d). Interestingly, all these mutations were located at the LC homodimer interface (Fig. 2e), suggesting that mutations in these positions may affect the structural integrity of the dimeric interface and/or induce local instability of the monomer, thus leading to LC misfolding and aggregation. A similar trend was also observed for other top-ranked features, where unmutated positions were, conversely, more frequent in *nox* sequences than in *tox* sequences (Fig. 2d).

To investigate the role of the top-ranked features in the prediction of LC toxicity, we performed a quantitative analysis of the importance of features identified by the feature selection technique. To this end, we trained 30 different classifiers (with a 10-fold cross validation) adding successively the 10 most important features of each feature family according to their information gain. Results are reported in Supplementary Figure 2 and Supplementary Table 9. Interestingly, the classifier using only the highest ranked feature 49A achieves an accuracy of 64% with an AUC of 0.55, while to achieve an AUC above 0.77 at least 17 top features are required.

Taken together, these findings show that the presence or absence of specific mutations at specific positions of an LC are key features used by LICTOR to classify the LC phenotype. More importantly, this further underlines the pivotal role of SMs as causative of AL.

### Reverting the toxic phenotype of an LC using LICTOR

Having assessed the prediction accuracy of LICTOR, we sought to validate the key toxicity determinants identified previously through information gain by computationally reverting the toxic phenotype of an LC with LICTOR and verifying the results in a validated *in vivo C. elegans* model^21,22^. We therefore selected an LC from our database (tox153) previously described in the literature as cardiotoxic^37^ and thus having the worst prognosis in AL. For this sequence, we had access to a bone marrow sample, from which we obtained the full-length sequence of tox153 (see Methods). Despite its toxic phenotype, tox153 differs by only 5 SMs from the corresponding germline (Fig. 3b), hence representing a good candidate with which to perform our study. From the analysis of the 5 SMs of tox153, we found that the best candidate, i.e., the mutation with the largest information gain, was at position 52, in which the germline leucine (LEU) was somatically mutated to a valine (VAL). This feature was one of the top-ranked predictor variables in all three families (AMP, MAP, and DAP). In fact, an unmutated amino acid at position 52 (52X) was significantly more frequent in *nox* sequences than in *tox* sequences (p_val_=1.5 e-09, Fig. 2d). Therefore, 52X may represent a “non-toxic feature” able to revert the phenotype of tox153 (Fig. 3c). To test this hypothesis, we restored the leucine of the germline sequence at position 52 of tox153 (tox153V52L) and used LICTOR to predict the toxicity of the new sequence. LICTOR predicted tox153V52L as a *nox* sequence, highlighting that this single point mutation is able to completely revert the toxic phenotype of tox153, according to *in silico* prediction. Next, we analysed the other 4 SMs according to their information gain. Among these SMs, only the MAP feature 56X-59X (p_val_ = 2.2 e-16, Fig. 2d) was included in the top ranked. Since tox153 is mutated at position 56 but not at position 59, we also reverted this SM in tox153V52L by mutating alanine to glycine (tox153V52LA56G) (Fig. 3c). Interestingly, *in silico* prediction by LICTOR confirmed the non-toxic phenotype of tox153V52LA56G. These results underline that key predictor variables identified in the feature selection process and used by LICTOR to perform the predictions represent molecular determinants of AL.

### Experimental validation of LICTOR using *C. elegans*

Next, we assessed the accuracy of LICTOR’s toxicity predictions in a validated *in vivo* model, exploiting the ability of *C. elegans* to specifically identify cardiotoxic LCs^21,22^. To this end, recombinantly expressed tox153, single mutant tox153V52L, double mutant tox153V52LA56G and tox153 germline protein (tox153GL), the protein without SMs, were administered to worms, and their toxicity was evaluated by measuring alterations in the pharyngeal pumping rate^21,22^. Tox153 caused significant pharyngeal dysfunction (Fig. 3d) comparable to that induced by the administration of cardiotoxic LCs purified from patients suffering from AL (H6, H7, H9) (Fig. 3e) ^21,22,38^, while the effect of tox153GL was comparable to that of the vehicle. The presence of a single mutation in tox153V52L significantly decreased the ability of the wild-type protein to cause pharyngeal toxicity (Fig. 3d). Notably, the double mutant tox153V52LA56G, similar to LCs purified from patients affected by multiple myeloma (M2, M7, M8)^21,22^, did not display toxic activity (Fig. 3e).

Finally, we thought to validate our starting hypothesis that LCs acquire toxic features trough the addition of specific SMs during the process of affinity maturation and that, conversely, germline LCs are never associated with AL development. To achieve this, we exploited the *C. elegans* model and tested a total of 3 germline LCs recombinantly expressed (H6GL, H9GL and Tox153GL). Values of their pharyngeal toxicity where then compared to those of the corresponding cardiotoxic LCs (H6, H9 and tox153) for which the pharyngeal bulb contraction (pumps/min) values were already published^21,22^ and obtained under the same experimental conditions, or measured by us (Figure 3d and Figure 3e). Interestingly, the three germline LCs do not show any significant proteotoxicity. Finally, we tested an additional cardiotoxic LC (H18) belonging to the same germline family of tox153 (Figure 3d), whose sequence was present in the *valset*. As expected H18 caused a significant impairment of *C. elegans* pharyngeal activity, while all germline proteins did not affect the pharyngeal pumping rate (Figure 3d and 3e).

Globally, these results experimentally validate our starting hypothesis that SMs are pivotal determinants of LC toxicity. Moreover, our *in vivo* analysis confirms the soundness of LICTOR prediction and the validity of key predictor variables identified by information gain as determinants of *in vivo* proteotoxicity.

## Discussion

Early diagnosis of AL is essential to readily apply therapeutic interventions and prevent permanent and fatal organ damage. However, AL is usually detected only once symptoms reflecting advanced organ involvement occur, which results in poor patient prognosis. Moreover, although pre-existing MGUS is a known risk factor for AL, predicting whether MGUS patients will progress to AL remains an open, unsolved problem. The extreme sequence diversity of LCs responsible for AL, due to VJ recombination and SMs, further complicates this scenario. Consequently, to deepen our understanding of the AL determinants and ultimately foster early AL diagnosis, we investigated LC sequences with a known clinical phenotype, with the aim of devising a predictive tool able to flag toxic LCs in advance (i.e., LCs responsible for the formation of toxic aggregates and AL development). To achieve this goal, we analysed a large dataset of toxic (*tox*) and non-toxic (*nox*) LCs of the λ isotype, the most frequent isotype in AL, following the hypothesis, also posed by other research groups^39–41^, that specific SMs can increase the propensity of LCs to cause AL. Therefore, we performed a statistical analysis of the distribution of SMs between *tox* and *nox* sequences. This analysis revealed that toxic LCs have significantly higher SM frequencies than non-toxic LCs (Fig. 1c). Based on these findings, we designed LICTOR, a machine-learning approach using SMs to classify the LC phenotype. LICTOR achieved a specificity and a sensitivity of 0.82 and 0.76, respectively, with an AUC of 0.87, making it an unprecedented tool in early AL diagnosis. Interestingly, including LC germline VJ rearrangements as additional predictor variables in LICTOR configuration did not improve prediction performance, further suggesting that, despite the prevalence of some VL germline genes in AL, SMs represent the critical driver of the disease.

LICTOR differs from approaches, such as AGGRESCAN^42^, PASTA^43^, WALTZ^44^ and others^45–48^, that predict the aggregation propensity of a protein by identifying amyloidogenic regions. LICTOR instead, aims at finding hotspots responsible for LC toxicity in AL amyloidosis, starting from the known clinical phenotype of LC sequences and following the assumption that SM are the key determinants of LC proteotoxicity. Indeed, our approach uses an innovative encoding scheme to express LC sequences as the difference with the respect to the germline. Then, this strategy is applied to extract sequence and structural features, and to investigate their role in the determination of LC proteotoxicity. Indeed, we showed that specific features (sequence AMP or structural MAP and DAP features) that provide the largest information gain to LICTOR harbour crucial information to accurately predict LC phenotype and can thus be regarded as effective AL molecular determinants. In fact, through the information gain feature selection process, we identified a set of features characterized by the strongest association with the LC phenotype, which, remarkably, were mainly located at the dimeric interface of the LC structure. This finding further emphasizes the key role of the structural context of SMs as drivers of LC proteotoxicity.

We also performed a comparison with AGGRESCAN, PASTA and WALTZ, using the aggregation propensities (P_A_ for AGGRESCAN, P_P_ for PASTA and P_w_ for WALTZ) provided by the respective programs to construct three classifiers. Interestingly, in all three cases *tox* sequences are significantly more aggregation prone than *nox* sequences (for AGGRESCAN mean P_A_ = −8.53±4.2 vs −7.44±3.8, p-val < 0.0001 unpaired t-test; for PASTA mean P_P_ = −6.35±1.08 vs −5.82 ± 1.15, p-val = < 0.0001 unpaired t-test; for WALTZ mean P_W_ = 97.78±0.83 vs 97.49±0.89, p-val =0.009 unpaired t-test). However, given the considerable overlap between the propensity distributions, classifiers based on aggregation propensity have a rather limited accuracy (AGGRESCAN accuracy = 0.59, PASTA accuracy = 0.68, WALTZ accuracy = 0.64). Nevertheless, these approaches indicate that toxic sequences are more prone to aggregate than non-toxic ones, a fact that is mirrored in LICTOR by the identification of SMs clustering at the dimer interface as divers of proteotoxicity in AL.

Previous studies have analyzed the fibril formation propensity of LC variable domains (VL), showing that destabilizing mutations at specific structural sites, correlate with increased amyloid fibril formation^49–51^. Additional reports have also compared the stability and fibril formation propensity of variable domains of toxic LC sequences associated with AL and non-toxic ones associated with multiple myeloma (MM)^52,53^, suggesting that specific mutations could induce a destabilization in toxic LCs. However, as pointed out in a recent analysis of full-length LCs associated with AL or MM^29^, despite significant differences in some properties such as the melting temperature (T_m_), it is not possible to unequivocally differentiate between pathogenic and non-pathogenic LCs based on a single biophysical property. Only flexibility and susceptibility to protease cleavage emerged as discriminative factor of proteotoxicity.

Other reports have suggested that germline proteins are more stable than the corresponding pathological LCs^39,54^. Analyzing the variable domains of the pathogenic light chain AL09 and its corresponding germline κI O18/O8, Baden et al., noticed that a non-conservative somatic mutation at the dimeric interface of AL09 (Y87H, according to Kabat numbering scheme), induced an altered dimer interface, characterized by a 90° rotation with respect to the canonical homodimeric structure of the germline counterpart. Interestingly, the same position (position 99 according to our sequential numbering scheme, corresponding to position 87 in Kabat-Chothia numbering) was also identified as one of the most important structural features (Figure 2d and Supplementary Table 8) used by LICTOR to predict LC toxicity.

CryoEM structures of LC amyloid fibrils^55,56^ show an interesting rearrangement in the region comprising of the intrachain disulfide bond. Namely, in folded LCs these two cysteines connect parallel ß-strands, while in the amyloid fibrils the two ß-strands are antiparallel. These conformational rearrangements break the intrachain interaction between CDR1 and CDR3, as well as, the intrachain interactions between FR2 and end of FR3. Furthermore, the dimerization interface of the folded LC is disrupted in the fibrils, as they are on the opposite side of the fibril layer. These findings are in line with our results that suggests that SM located at the LC homodimeric interface may impair the structural integrity of the protein-protein interface and/or induce local instability of the monomer, with consequent triggering of LC misfolding and the generation of toxic species.

The starting hypothesis that SMs are key determinants of AL, the accuracy of LICTOR and of our *in silico* findings were also experimentally confirmed in *C. elegans*, a validated *in vivo* model for assessing LC toxicity. We demonstrated that germline LCs are not able to induce proteotoxicity *in vivo*, validating the assumption that naïve LC sequences acquire the toxic phenotype in AL during affinity maturation. Notably, as predicted by LICTOR, the toxic phenotype of an LC was abolished by reverting a single SM.

Taken together, these findings confirm the accuracy and robustness of our *in silico* approach in the identification of toxic and non-toxic LCs and suggest its usefulness as a diagnostic instrument for AL.

Machine learning relies on data. Larger datasets of LC sequences would, therefore, be beneficial for validation, as well as for the improvement of LICTOR accuracy. However, collecting large numbers of toxic LC sequences is difficult due to the low prevalence of the disease. We believe that the application of LICTOR as a possible diagnostic tool could encourage clinicians to obtain – and make available to the public – LC sequences of AL patients, thus increasing the size of LC sequence databases and consequently allowing improvement of LICTOR accuracy. Furthermore, other factors such as the increased protein dynamics of toxic LCs^29^ or the generation of LC glycosylation sites by somatic mutations, may be included among the predictive features to improve LC toxicity classification, as suggested by previous reports^51,57^. This would not only improve the accuracy of LICTOR, but also deepen our understanding of AL determinants and shed light on the complex mechanism of AL development.

In conclusion, LICTOR represents the first method for the accurate prediction LC toxicity from their sequence, allowing the timely identification of high-risk patients, such as MGUS subjects likely to progress to AL. Using LICTOR can thus promote a closer monitoring for AL development and foster early treatment and better patients’ prognosis. Finally, LICTOR may be used together with other recently proposed strategies, such as the differential recruitment efficacy of patients derived full-length LCs by synthetic amyloid fibrils^58^, to predict the risk of AL development. Our approach may, furthermore, guide the development of novel predictive tools useful for other diseases, such as cancer, in which the prognosis may depend on SMs of specific tumour-linked proteins. LICTOR is available as a webservice at http://lictor.irb.usi.ch.

## Methods

### Dataset

The database used in the training was composed of 428 *tox* and 590 *nox* sequences of the λ isotype collected from the Amyloid Light-chain Database (ALBase) (http://albase.bumc.bu.edu). Furthermore, it contained 57 *nox* λ LC sequences that we collected at the Institute for Research in Biomedicine (IRB-DB), known to be non-toxic in the context of AL. The 1,075 sequences were automatically aligned using a progressive Kabat-Chothia numbering scheme (http://www.bioinf.org.uk/abs/). According to this scheme, for example, the CDR1 of a given LC with Kabat-Chothia numbering 30A, 30B, 30C, 30D, 30E, and 30F was assigned 31, 32, 33, 34, 35, and 36. For the ALBase sequences, germline information was taken from the database, while for IRB-DB LCs, the germline was assessed with an in-house script. Next, germline (GL) sequences were reconstructed using the IMGT database^25^.

The GL sequences were aligned with the same numbering scheme used for the LCs. Next, each LC in the dataset was compared with the corresponding GL to identify all somatic mutations, with the differences encoded using an X for unmutated positions and the LC amino acid for somatic mutations; this sequence was referred to as S_*mut*_. For example, an LC with sequence *SYELTQPP* and a corresponding GL with the sequence *SYVLTQPP* was encoded as *XXEXXXXX* since there is a somatic mutation (*V*→*E*) at position 3. To compare the presence of somatic mutations in S_*mut*_ at each position *i* in the Kabat-Chothia numbering scheme, the following four quantities were computed:

- 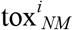- the number of toxic sequences without a somatic mutation at position *i;*
- 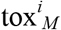- the number of toxic sequences with a somatic mutation at position *i;*
- 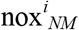- the number of non-toxic sequences without a somatic mutation at position *i;*
- 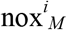- the number of non-toxic sequences with a somatic mutation at position *i.*

### Statistical analysis

The *fisher.test* function in R version 3.5.1 with the arguments *conf.int*=*TRUE* and *conf.level*=0.95 was used to assess significant differences in somatic mutations between toxic and non-toxic sequences. The OR between 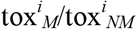 and 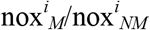 was computed as

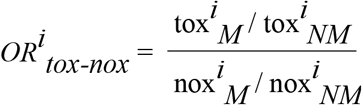

OR=1 indicates that the event under study (i.e., the frequency of mutations at position *i*) is equally likely in the two groups (e.g., *tox* vs *nox*). OR>1 indicates that the event is more likely in the first group (*tox*). OR<1 indicates that the event is more likely in the second group (*nox*). The t.test function in R version 3.5.1 was used to evaluate whether the probability distributions of the number of SMs (PDSM) differed between the *tox*, *nox* and *hdnox* datasets. Data from *C. elegans*-based assays were analyzed using GraphPad Prism 8.2.1 software by one-way ANOVA and Dunn’s post-test analysis. A *P value* < 0.05 was considered significant.

### Predictor variables used by the machine learners

Given a sequence, the following features were extracted:

#### Amino acid at each Mutated Position (AMP)

From a sequence *S_mut_*, a list of predictor variables was extracted, each describing the type of amino acid added by the somatic mutation at a given position or the absence of a mutation at the position. Thus, each of these variables was a pair (*position*, *amino acid*), where we used the letter “X” instead of the amino acid at the positions for which no somatic mutations were present.

#### Monomeric Amino acid Pairs (MAP)

LCs share a conserved 3D structure. Therefore, pairs of interacting residues were defined as amino acids having a distance between the respective Cβ atoms less than 7.5 Å in the X-ray structure (PDB ID: 2OLD).

#### Dimeric Amino acid Pairs (DAP)

Similarly, pairs of residues that interact at the LC-LC interface were defined using the 2OLD LC homodimeric X-ray structure. Two residues belonging to different chains were considered to interact if the distance between their Cβ atoms was less than 7.5 Å.

### Machine learning algorithms

Weka 3.8.1^30^ implementation was used for the four machine learning algorithms (Bayesian network, logistic regression, J48, and random forest) to solve the classification task. For all algorithms, the default Weka parameters were used. The algorithms were evaluated by performing 10-fold cross-validation over the dataset. The performance of each algorithm was first assessed using only one family of features (e.g., AMP, MAP, and DAP, for a total of three combinations); second, the three families were combined into pairs (e.g., AMP U MAP, for a total of three combinations); third, all three families were combined together. This led to a total of 7 (feature configurations) × 4 (algorithms) = 28 prediction experiments. Moreover, each of the 28 experiments was performed with and without the balancing of the training set with SMOTE (**S**ynthetic **M**inority **O**ver-sampling **TE**chnique)^31^ on the toxic sequences so that the number of toxic instances was equal to the number of non-toxic instances in the training set during each of the ten cross-validations used in the evaluation. This led to 28 × 2 (with/without SMOTE) = 56 total experiments.

### Prediction performance

The various prediction algorithms were assessed by computing the following classification errors: (i) Type I misclassifications, indicating toxic sequences incorrectly classified as non-toxic (false negative—FN), and (ii) Type-II misclassifications, indicating non-toxic sequences misclassified as toxic (false positive—FP). The correct classifications were instead indicated by the number of true positives—TP (a toxic sequence correctly classified) —and true negatives—TN (a non-toxic sequence correctly classified).

Based on TP, TN, FP, and FN, the following metrics were used to evaluate the performance of our classifiers:

- *Area under the receiver operating characteristic curve (AUC).* The AUC is used to assess the performance of a two-class classifier (such as that in our study) and is equal to the probability that the classifier will rank a randomly chosen positive instance (in our case, a toxic sequence) higher than a randomly chosen negative instance (a non-toxic sequence). A random classifier has an AUC=0.5, while the AUC is 1.0 for a perfect classifier.
- *Sensitivity.* Computed as TP/(TP+FN), this represents the percentage of toxic sequences correctly identified by the classifier.
- *Specificity.* Computed as TN/(TN+FP), this represents the percentage of non-toxic sequences correctly identified by the classifier.
- *Accuracy.* Computed as (TP+TN)/(TP+FP+TN+FN), this represents the overall percentage of correctly classified sequences.
- *Balanced accuracy*. Computed as (Specificity+Sensitivity)/2, this represents the arithmetic mean of Sensitivity and Specificity.
- *F1* score. Computed as 2TP/(2TP+FP+FN), this represents the harmonic mean of the Sensitivity and the Precision, which is computed as the number of TP/(TP+FN). F1 = 1 indicates perfect Precision and Sensitivity, while F1 = 0 represents the lowest possible value achieved if either the Precision or the Sensitivity is 0.

### Youden index

The Youden (J) index was used to validate the effectiveness of the predictors and to find the optimal cut-off point to separate toxic LCs associated with the disease from non-toxic LCs using the following formula:

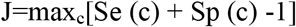

### Information gain feature selection

The InfoGainAttributeEval filter implemented in Weka 3.8.1^36^ was used to remove all features that did not contribute to the information available for the prediction of the sequence type. All features having an information gain less than 0.01 were removed. Given the computational cost of this procedure, this experiment was performed for the best-performing algorithm and configuration identified in the previous 56 experiments. The full list of ranked features is shown in Supplementary Table 8.

### Tox153 sequence

The full-length LC sequence was obtained from bone marrow cells with TRIzol lysate. RNA was extracted using Direct-zol RNA Miniprep Zymo-Spin IIC columns (Zymo Research) and used for cDNA synthesis, employing the template switching technique approach^59^. Ig genes were amplified with specific primers, and a library of unique sequences was obtained by a Zero Blunt® TOPO® PCR Cloning Kit (Life Technologies) and subsequent analysis of single colonies.

### Protein production and purification

Tox153, tox153V52L and tox153V52LA56G were custom expressed in mammalian cell lines (Expi293F), purified by affinity purification column, and analysed by SDS-Page and Western blot by GenScript (New Jersey, USA). H6GL, H9GL, Tox153GL and H18 were expressed in mammalian cell lines (Expi293F), purified by HiTrap® LambdaFabSelect (GE Healthcare), and analysed by SDS-Page.

### Effect of LCs on *C. elegans*

Bristol N2 nematodes were obtained from the *Caenorhabditis elegans* Genetic Center (CGC, University of Minnesota, Minneapolis, MN) and propagated at 20°C on solid nematode growth medium (NGM) seeded with *Escherichia coli* OP50 (CGC) for food. Worms were incubated with 100 µg/ml tox153 wild-type protein, tox153V52L or tox153V52LA56G (100 worms/100 µl) in 10 mM phosphate-buffered saline (PBS, pH 7.4)^8,9^. Hydrogen peroxide (1 mM) was administered under dark conditions as a positive control and 10 mM PBS (pH 7.4) as a negative control (vehicle). After 2 h of incubation with orbital shaking, worms were transferred onto NGM plates seeded with OP50 *E. coli*. The pharyngeal pumping rate, measured by counting the number of times the terminal bulb of the pharynx contracted over a 1-min interval, was scored 24 h later.

## Acknowledgements

This study was supported by grants from the Swiss National Science Foundation (31003A-166472) to A.C. and from the Italian Ministry of Health (RF-2013-02355259 and RF-2016-02361756) to L.D. *C. elegans* and OP50 *E. coli* were provided by the CGC, which is funded by NIH Office Research Infrastructure Programs (P40 OD010440). We would like to acknowledge the use of the Boston University ALBase, supported by HL68705, and Dr. Giampaolo Merlini and Dr. Mario Nuvolone, who provided bone marrow samples.

## Author contributions

A.C. and M.G. designed the research. M.G. performed data acquisition and analysis and drafted the manuscript. M.F., L.P., S.R., M.R. M.M.B, R.D.G., L.V., L.D. and A.L. provided data. M.G., L.P., S.R., M.F., M.M., J.S., R.D.G., M.P., O.M., A.L., L.D. and A.C. edited and approved the manuscript.

