## Supplementary Figures for "Machine learning predicts immunoglobulin light chain toxicity through somatic mutations"

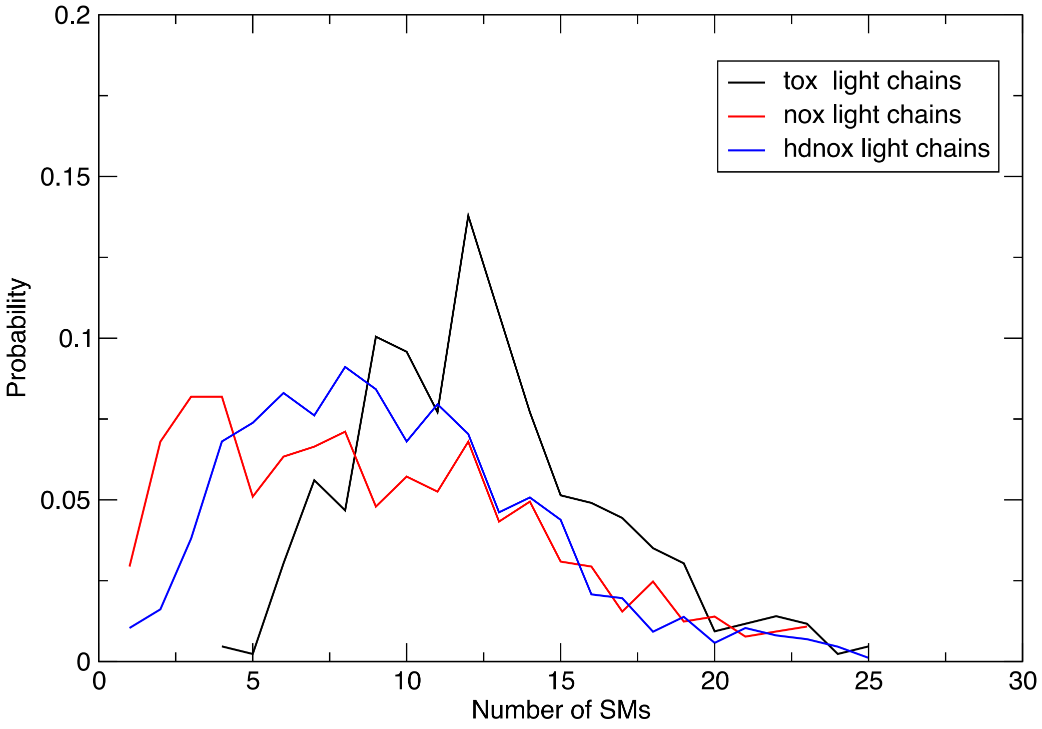


**Supplementary Figure. 1 | Probability distribution of somatic mutations in toxic and non-toxic light chain sequences.** The black line represent the probability distributions of somatic mutations (PDSM) in toxic light chains (tox) (average number of mutations equal to 12.5), the red line represent the PDSM of non-toxic sequences (nox) used in the training of the machine learning algorithm (average number of mutations equal to 8.8), while the blue line represent the PDSM computed randomly selecting 1000 lambda LCs from a healthy donor[1] (hdnox) (average number of mutations equal to 9.6). The difference in average number of somatic mutations in the two set of non-toxic LC sequences is not statistically significant ($p = 0.1$), while there is a statistically significant difference between the toxic sequences and both non-toxic dataset ($p < 10e-29$). Statistical analysis were performed using Student *t* test.

**a**

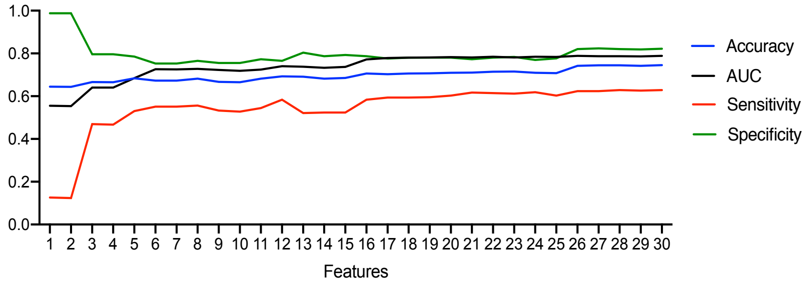


**b**

| | Features |
|---|---|
| 1 | 49A |
| 2 | 49A.I49AX116I |
| 3 | 49A.I49AX116I.I65XX107I |
| 4 | 49A.I49AX116I.I65XX107I.I4XA49I |
| 5 | 49A.I49AX116I.I65XX107I.I4XA49I.I49XX99I |
| 6 | 49A.I49AX116I.I65XX107I.I4XA49I.I49XX99I.I52XX108I |
| 7 | 49A.I49AX116I.I65XX107I.I4XA49I.I49XX99I.I52XX108I.I49AX99I |
| 8 | 49A.I49AX116I.I65XX107I.I4XA49I.I49XX99I.I52XX108I.I49AX99I.107X |
| 9 | 49A.I49AX116I.I65XX107I.I4XA49I.I49XX99I.I52XX108I.I49AX99I.107X._104XX108_ |
| 10 | 49A.I49AX116I.I65XX107I.I4XA49I.I49XX99I.I52XX108I.I49AX99I.107X._104XX108_.I3XA49I |
| 11 | 49A.I49AX116I.I65XX107I.I4XA49I.I49XX99I.I52XX108I.I49AX99I.107X._104XX108_.I3XA49I.106X |
| 12 | 49A.I49AX116I.I65XX107I.I4XA49I.I49XX99I.I52XX108I.I49AX99I.107X._104XX108_.I3XA49I.106X._44XX98_ |
| 13 | 49A.I49AX116I.I65XX107I.I4XA49I.I49XX99I.I52XX108I.I49AX99I.107X._104XX108_.I3XA49I.106X._44XX98_._52XX65_ |
| 14 | 49A.I49AX116I.I65XX107I.I4XA49I.I49XX99I.I52XX108I.I49AX99I.107X._104XX108_.I3XA49I.106X._44XX98_._52XX65_._44XX99_ |
| 15 | 49A.I49AX116I.I65XX107I.I4XA49I.I49XX99I.I52XX108I.I49AX99I.107X._104XX108_.I3XA49I.106X._44XX98_._52XX65_._44XX99_.I66XX107I |
| 16 | 49A.I49AX116I.I65XX107I.I4XA49I.I49XX99I.I52XX108I.I49AX99I.107X._104XX108_.I3XA49I.106X._44XX98_._52XX65_._44XX99_.I66XX107I._54XX58_ |
| 17 | 49A.I49AX116I.I65XX107I.I4XA49I.I49XX99I.I52XX108I.I49AX99I.107X._104XX108_.I3XA49I.106X._44XX98_._52XX65_._44XX99_.I66XX107I._54XX58_._1XH108_ |
| 18 | 49A.I49AX116I.I65XX107I.I4XA49I.I49XX99I.I52XX108I.I49AX99I.107X._104XX108_.I3XA49I.106X._44XX98_._52XX65_._44XX99_.I66XX107I._54XX58_._1XH108_.I49XX116I |
| 19 | 49A.I49AX116I.I65XX107I.I4XA49I.I49XX99I.I52XX108I.I49AX99I.107X._104XX108_.I3XA49I.106X._44XX98_._52XX65_._44XX99_.I66XX107I._54XX58_._1XH108_.I49XX116I.108H |
| 20 | 49A.I49AX116I.I65XX107I.I4XA49I.I49XX99I.I52XX108I.I49AX99I.107X._104XX108_.I3XA49I.106X._44XX98_._52XX65_._44XX99_.I66XX107I._54XX58_._1XH108_.I49XX116I.108H._44HX96_ |
| 21 | 49A.I49AX116I.I65XX107I.I4XA49I.I49XX99I.I52XX108I.I49AX99I.107X._104XX108_.I3XA49I.106X._44XX98_._52XX65_._44XX99_.I66XX107I._54XX58_._1XH108_.I49XX116I.108H._44HX96_._44XX97_ |
| 22 | 49A.I49AX116I.I65XX107I.I4XA49I.I49XX99I.I52XX108I.I49AX99I.107X._104XX108_.I3XA49I.106X._44XX98_._52XX65_._44XX99_.I66XX107I._54XX58_._1XH108_.I49XX116I.108H._44HX96_._44XX97_.78X |
| 23 | 49A.I49AX116I.I65XX107I.I4XA49I.I49XX99I.I52XX108I.I49AX99I.107X._104XX108_.I3XA49I.106X._44XX98_._52XX65_._44XX99_.I66XX107I._54XX58_._1XH108_.I49XX116I.108H._44HX96_._44XX97_.78X.I52XX116I |
| 24 | 49A.I49AX116I.I65XX107I.I4XA49I.I49XX99I.I52XX108I.I49AX99I.107X._104XX108_.I3XA49I.106X._44XX98_._52XX65_._44XX99_.I66XX107I._54XX58_._1XH108_.I49XX116I.108H._44HX96_._44XX97_.78X.I52XX116I.106N |
| 25 | 49A.I49AX116I.I65XX107I.I4XA49I.I49XX99I.I52XX108I.I49AX99I.107X._104XX108_.I3XA49I.106X._44XX98_._52XX65_._44XX99_.I66XX107I._54XX58_._1XH108_.I49XX116I.108H._44HX96_._44XX97_.78X.I52XX116I.106N._42XX52_ |
| 26 | 49A.I49AX116I.I65XX107I.I4XA49I.I49XX99I.I52XX108I.I49AX99I.107X._104XX108_.I3XA49I.106X._44XX98_._52XX65_._44XX99_.I66XX107I._54XX58_._1XH108_.I49XX116I.108H._44HX96_._44XX97_.78X.I52XX116I.106N._42XX52_._56XX59_ |
| 27 | 49A.I49AX116I.I65XX107I.I4XA49I.I49XX99I.I52XX108I.I49AX99I.107X._104XX108_.I3XA49I.106X._44XX98_._52XX65_._44XX99_.I66XX107I._54XX58_._1XH108_.I49XX116I.108H._44HX96_._44XX97_.78X.I52XX116I.106N._42XX52_._56XX59_.49X |
| 28 | 49A.I49AX116I.I65XX107I.I4XA49I.I49XX99I.I52XX108I.I49AX99I.107X._104XX108_.I3XA49I.106X._44XX98_._52XX65_._44XX99_.I66XX107I._54XX58_._1XH108_.I49XX116I.108H._44HX96_._44XX97_.78X.I52XX116I.106N._42XX52_._56XX59_.49X.44X |
| 29 | 49A.I49AX116I.I65XX107I.I4XA49I.I49XX99I.I52XX108I.I49AX99I.107X._104XX108_.I3XA49I.106X._44XX98_._52XX65_._44XX99_.I66XX107I._54XX58_._1XH108_.I49XX116I.108H._44HX96_._44XX97_.78X.I52XX116I.106N._42XX52_._56XX59_.49X.44X.52X |
| 30 | 49A.I49AX116I.I65XX107I.I4XA49I.I49XX99I.I52XX108I.I49AX99I.107X._104XX108_.I3XA49I.106X._44XX98_._52XX65_._44XX99_.I66XX107I._54XX58_._1XH108_.I49XX116I.108H._44HX96_._44XX97_.78X.I52XX116I.106N._42XX52_._56XX59_.49X.44X.52X.44H |

**Supplementary Figure. 2 | Quantitative analysis of the importance of features identified by the feature selection technique training 30 different classifiers with a 10-fold cross validation, adding one by one the 10 most important features of each feature family according to information gain.** **a,** the contribution of each feature in the achieved AUC, Accuracy, Sensitivity and Specificity, and **b**, the specific features added. The results of these analysis are reported in Supplementary Table 9.
