## Supplementary Tables information for "Machine learning predicts immunoglobulin light chain toxicity through somatic mutations"

The Supplementary Tables n. 1, 3, 4, 5, 6, 7 and 9 included in the supplementary files report, for each of the ten folds used as test set, the results achieved in terms of True Positive (TP), True Negative (TN), False Positive (FP), and False Negative (FN). Based on this data, values for Sensitivity, Specificity, Accuracy, Balanced Accuracy, AUC and F1 score are reported as well. To see the overall results of each classifier (i.e., the results obtained across the ten folds), select “overall” in the Fold column.

**Supplementary Table 1 |** Results of the 56 experiments run with different combinations of predictor variables for the four machine learning algorithms.

The Random Forest configuration using AMP, MAP, and DAP as predictor variables is the one implemented in LICTOR.

**Supplementary Table 2 |** LC sequences not present in the training set and used to test LICTOR. The toxic LC sequences were derived from patients affected by AL with cardiac involvement (H3, H6, H7, H9, H15, H16, H18) and the non-toxic ones derived from patients affected by multiple myeloma (M2, M7, M8, M9, M10). All these LC were previously published[2].

**Supplementary Table 3 |** Results of LICTOR obtained when classifying 100 non-toxic light chain sequences randomly selected form a healthy donor repertoire (*hdnox*).

**Supplementary Table 4 |** Results obtained when classifying *tox* sequences randomly assigned to the *nox* or to the *tox* family.

**Supplementary Table 5 |** Results obtained when classifying *nox* sequences randomly assigned to the *nox* or to the *tox* family.

**Supplementary Table 6 |** Results of the *germline-based* predictors for the four machine learners.

**Supplementary Table 7 |** Results of LICTOR when using also LC germline VJ rearrangement as predictor variable.

**Supplementary Table 8 |** Results of the feature selection process.

The first column (average merit), reports the average information gain obtained when experimenting across the ten folds. The second column (average rank) reports the average rank of the feature across the ten folds (the higher the rank, the more important is the feature, with the most important feature in position 1). Finally, the third column (attribute) show a progressive id representing the position of the feature, followed by the feature itself. Features belonging to the DAP family are reported between "|" symbols. Those part of the MAP family between "_" symbols. The absence of both symbols indicates features belonging to the AMP family.

**Supplementary Table 9 |** Results obtained when training 30 different classifiers, with a 10-fold cross validation, adding one by one the 10 most important features of each feature family according to information gain.


1 DeKosky, B. J. *et al.* Large-scale sequence and structural comparisons of human naive and antigen-experienced antibody repertoires. *Proc Natl Acad Sci U S A* **113**, E2636-2645, doi:10.1073/pnas.1525510113 (2016).

2 Oberti, L. *et al.* Concurrent structural and biophysical traits link with immunoglobulin light chains amyloid propensity. *Sci Rep* **7**, 16809, doi:10.1038/s41598-017-16953-7 (2017).
